# Proteomes reflect size-based growth constraints across diverse microbes

**DOI:** 10.64898/2026.10.06.756872

**Authors:** J. Scott P. McCain, Teemu P. Miettinen, Rogier Braakman, Michael J. Follows

## Abstract

The maximum growth rate of microbes plays a central role in shaping ecological outcomes and is a major target of bioengineering efforts. Previous data compilations suggest single cell maximum growth rates mostly decrease with cell volume across species, except for an initial increase from the very smallest microbes to an intermediate size. Here we hypothesized that this unimodal relationship is shaped by the volumetric requirements of ribosomes and the surface area-dependence of nutrient supply. We developed a mechanistic model relating ribosomal protein mass fraction to growth rate and cell size. Predictions from this model were empirically supported by proteomic data from 97 prokaryotes ranging over 3 orders of magnitude in cell volume. Our analysis suggests that the smallest cells’ growth rates are restricted by the volumetric constraints on ribosomes per cell and rely on compact, heterotrophic metabolisms. In contrast, growth rates of the largest prokaryotes face geometric constraints on specific resource acquisition rates, capping growth rates and investment in ribosomal protein, but allowing investment in protein-expensive metabolisms including phototrophy and sulfur oxidation. Prokaryotic cells of intermediate size face neither constraint and exhibit a higher maximum growth rate potential, which, we hypothesize, is associated with elevated metabolic diversity.

## 1 Introduction

Maximum population growth rate is a key trait that shapes microbial communities under both high and low resource concentrations (hereafter just ‘maximum growth rate’). MacArthur and Wilson (1967) posited that high resource conditions select for fast growth rates. Under low nutrient conditions, maximum growth rate is a factor in determining the degree to which nutrients are drawn down, and therefore predictive of competitive outcomes (Tilman 1982, 1977). Maximum growth rate is also a central parameter in diverse ecosystem models, from those of the human gut to leaf litter to the global ocean (Smith et al. 2020; Allison 2012; Coles et al. 2017; Follows et al. 2007). Microbial bioengineering efforts often target achieving the fastest possible biomass accumulation, highlighting a practical importance of quantifying the biophysical limits on maximum growth rate. Despite the widespread importance of maximum growth rate, the exact mechanisms that underpin an organism’s maximum growth rate are uncertain.

There is a striking relationship between maximum growth rate (1/*time*) and cell size across single-celled microbes. The maximum growth rate increases from the very smallest prokaryotes (*<* 0.1 *µm*^3^) until a size of *∼*1-10 *µm*^3^, after which it declines in both prokaryotes and eukaryotes (Raven 1994; Marañón et al. 2013; Westoby et al. 2021; Lynch et al. 2022; Trickovic and Lynch 2025). This unimodal pattern can be seen another way – there is a single, fastest growing microbe: *Vibrio natriegens* (Weinstock et al. 2016). By definition, maximum growth rate must decrease for organisms smaller or larger than *V. natriegens*. Several models have been put forward to explain parts of this pattern. Low maximum growth rates in the smallest cells are hypothesized to reflect the role of non-scalable components: Raven (1994) argued that DNA and cell-membranes limit the available volume for biosynthetic machinery in the smallest photosynthetic cells. Trickovic and Lynch (2025) argued that the increase in growth rate with cell size reflects a decreasing relative investment in lipid biosynthesis proteins, as lipids become a smaller proportion of total biomass, increasing the available investment for ribosomal proteins. The decreasing relationship between size and maximum growth rate at cell volumes greater than *∼* 10*µm*^3^ has been attributed to geometric constraints on nutrient acquisition related to surface area-to-volume changes in phytoplankton (Edwards et al. 2012; Aksnes and Egge 1991), long within-cell diffusion times (Schulz and Jørgensen 2001), or population genetic mechanisms (Lynch et al. 2022).

Here we present a single, mechanistic framework that explains the links between cell size and maximum growth rate across a wide microbial size spectrum. Our model is rooted in several concepts mentioned above: at smaller sizes, microbial maximum growth rates are constrained by non-scalable components and the available space for biosynthetic machinery, while larger microbes are constrained by limits on specific nutrient uptake rates related to the surface area-to-volume ratio (SA/V). These concepts are described by a mathematical model, which is formulated in terms of the ribosomal protein mass fraction and then tested against a compilation of proteomic data from diverse microbes. Finally, we explore the consequences of this model in the size regimes where we predict the strongest effects: the giant and ultrasmall prokaryotes. Overall, our results suggest that two simple physical constraints give rise to the relationship between the maximum growth rate and the size of the organism.

## 2 Results and Discussion

### 2.1 A model of two biophysical constraints on maximum growth rate

We formulate a model relating cell volume and maximum growth rate in terms of ribosomes, which are a dominant component of the total mass of biosynthetic machinery (Scott et al. 2010) and this formulation will provide a quantitative test through proteomic measurements (see below). Seminal work identified a linear relationship between the ribosomal protein mass fraction and growth rate in *Escherichia coli* and other model organisms (Schaechter et al. 1958; Molenaar et al. 2009; Scott et al. 2010). Following this work, we represent the growth rate (*µ*),

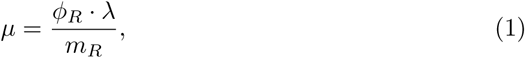

as a function of the relative investment in ribosomal proteins (ribosomal protein mass fraction; *ϕ_R_*, dimensionless), the number of amino acids per ribosome particle (*m_R_*), and the translation elongation rate (*λ*, amino acids *s*^-1^ ribosome^-1^). The ribosomal protein mass fraction is the ribosomal protein mass divided by the total protein mass per cell. Over evolutionary time scales, the investment in ribosomes, the ribosomal protein mass, or the elongation rate could be modulated to maximize growth rate: here we focus on ribosomal protein mass fraction (i.e., investment) as a function of organism size because it has the most observational constraints and exhibits significant variation.

We explore the relationship between maximum growth rate, cell volume, and ribosomal mass fraction as a consequence of two overarching physical constraints: i) For smaller cells, which have high nutrient affinities due to a high SA/V, non-scalable components limit the space for ribosomes and maximum growth rate is ribosome limited (Raven 1994). ii) For larger cells, which have no space constraint for ribosomes, low nutrient affinities due to low S/A ultimately limit their maximum growth rate (Fig. 1A) (Edwards et al. 2012).

**Fig. 1.**
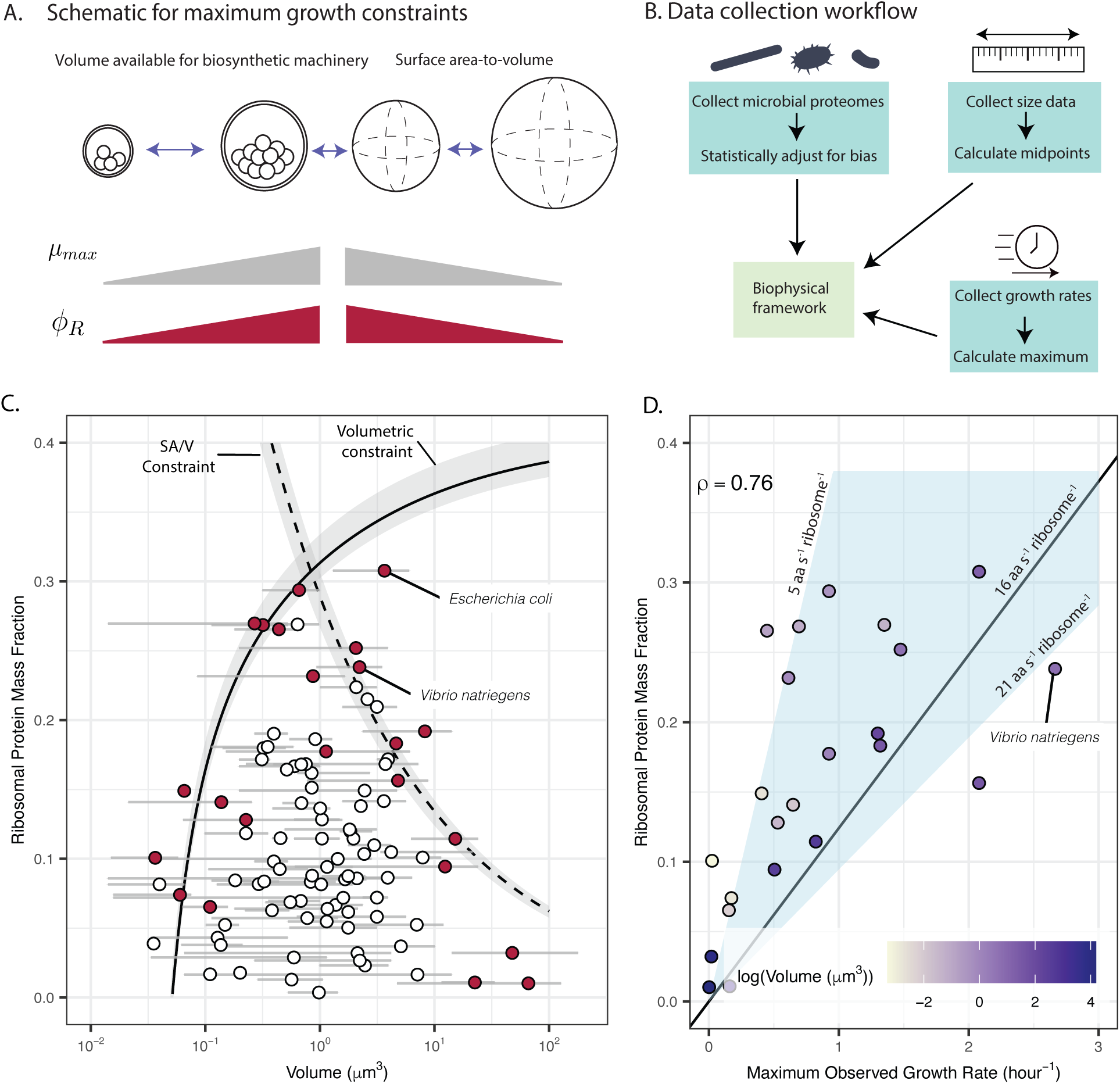
Size-based constraints on maximum growth rate in prokaryotes. (A) Hypothesized constraints on maximum growth rate: 1) the volume for biosynthetic machinery, which increases with cell volume, 2) the surface area-to-volume ratios of cells constrains nutrient supply rates and decreases with organism volume. These changes in maximum growth rate (*µmax*) are hypothesized to have an increasing (Equation (2)) and then decreasing (Equation (4)) pattern, which is accompanied by the same pattern in ribosomal protein mass fraction (*ϕ_R_*). (B) Schematic of the data collection workflow. First, proteomic data from various sources were used, and then ribosomal protein mass fraction was statistically estimated by adjusting for number of proteins detected (see Methods). Second, organism size and shape data were aggregated using published microscopy images and species descriptions. Finally, growth rate data were aggregated and the_4_ maximum for a given species was taken. (C) The ribosomal protein mass fraction across 97 prokaryotes as a function of volume. Each point corresponds to a unique species, specifically the maximum ribosomal protein mass fraction for that species. Error bars on volumes correspond to the maximum and minimum volumes observed for a given species, and the points are the midpoint volumes. Points in red were identified as part of the boundary of ribosomal protein mass fraction (see Methods for procedure to identify the boundary). The solid line corresponds to the fitted model prediction for the volumetric constraint, and the dashed line corresponds to the fitted model prediction for the surface area-to-volume constraint. Fitted parameter values were *ϕ_Q_* = 0.59 (dimensionless) and *kcat* = 82 (1/*s*). Grey regions surrounding both predictions are the minima and maxima when using variable numbers of bins for identifying ribosomal protein mass fraction maxima (see Methods). (D) The maximum observed growth rate as a function of ribosomal protein mass fraction for the red points from (C). Note that the black line in (D) is Equation 1 (i.e., not a fitted model), and it varies as a function of the assumed ribosomal elongation rate, which is shown next to the black line. The blue region surrounding the black line corresponds to different ribosomal elongation rates, also shown next to their associated lines. The Spearman correlation coefficient is displayed. The point colour corresponds to the natural logarithm of the midpoint volume for each organism.

#### (i) Smaller cells limited by space for ribosomes

For the smallest cells, the non-scalable component hypothesis (Raven 1994) argues that sufficient space for biosynthetic machinery constrains maximum growth rate. Here we explicitly focus on ribosomes as universal biosynthetic machinery. For very small prokaryotes, periplasmic space can occupy even 20-55% of total volume (Zhao et al. 2017; Luef et al. 2015), and some cellular components, such as DNA, have a minimum space requirement (Kempes et al. 2016; Raven 1994). Thus, volume available for ribosome content caps maximum growth rates in the smallest cells. As cell size increases, this volumetric constraint is relieved, so ribosomal protein mass fraction and maximum growth rate can also increase (Fig. 1A).

To model the volumetric constraints on ribosomes per cell, we consider the maximum potential ribosomal investment, describing the ribosomal protein mass fraction, 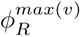 (dimensionless), as a function of the inner volume of a cell, *V_in_*, which is the total cell volume less that occupied by the envelope (*V_e_*; we assume constant envelope thickness across species). Ribosomes are restricted to the inner volume. We also account for the volume of DNA as a component of the inner volume (*C*; *µm*^3^) and a constant proportion of total protein is assumed to be required for other cellular processes like maintenance (*ϕ_Q_*; dimensionless; hereafter we refer to this as the maintenance proteome). The upper-bound on ribosomal investment, 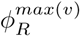 (where *max*(*v*) denotes the volumetric constraint), is therefore

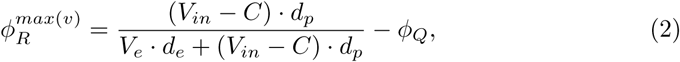

where *d_p_* and *d_e_* are the densities of cytosolic and envelope protein, respectively (amino acids *µm^−^*^3^). 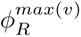 increases monotonically and saturates with cell size (Fig. 1C; solid line; saturating to a value of 1 *− ϕ_Q_*) as the space constraint on ribosomal investment weakens. We examine the upper bound on ribosomal protein mass fraction (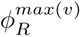; Equation 2) using inner (i.e., cytosolic) and envelope protein densities that have been quantified in model organisms (Chure et al. 2025; Cayley et al. 1991; Dai et al. 2016; Reuveni et al. 2017; Li and Du 2014; Menden-Deuer and Lessard 2000; see Supplementary Table S1 for all parameter values, and Supplementary Fig. S1 for a diagram of all parameters and their corresponding physical representation). The only fitted parameter in Equation 2 is *ϕ_Q_*, and we describe the fitting procedure and required data below. We note that this is a model of maximum ribosomal protein fraction: cells may invest in different allocation strategies that could reduce ribosomal content and growth rate. Assuming that the translation elongation rate, *λ*, and the amino acid content of ribosomes, *m_R_*, are constant, the maximum growth rate in this regime is linearly proportional to the predicted constrained 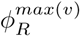 (Equation 1).

#### (ii) Larger cells limited by low nutrient affinities

We hypothesized that as larger cells are relieved of the volumetric constraint described above, their maximum potential growth rate is, instead, constrained by the specific rate of acquisition of external resources. The growth rate, *µ*, is the ratio of nutrient uptake to nutrient quotas, *µ* = *V_n_/Q_n_*, where *V_n_* is the cell-specific uptake rate (moles cell*^−^*^1^ *s^−^*^1^) and *Q_n_* is the cell-specific nutrient quota (moles cell*^−^*^1^). Maximum cell specific uptake rate is related to surface area available for transporters (Edwards et al. 2012; Zwanzig 1990; Berg and Purcell 1977) and cell quota increases with cell volume (Menden-Deuer and Lessard 2000). Under nutrient replete concentrations, nutrient uptake is maximized (*V_n_*→*V_max_*) and so is growth rate (*µ*→*µ_max_*). Assuming a spherical organism,

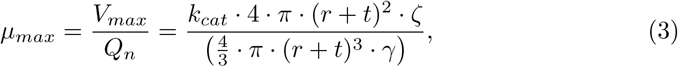

where *k_cat_* (*s^−^*^1^) is the maximum turnover rate per transporter, *γ* is the cellular nutrient concentration (moles *µm^−^*^3^; inclusive of nitrogen atoms that are a component of larger molecules), *r* is the inner cell radius (*µm*), and *t* is the envelope thickness (*µm*). The number of nutrient transporters per cell is constrained by the surface area, and the nutrient transporter density on the surface of the cell is *ζ* (transporters *µm^−^*^2^). Therefore, the maximum growth rate is inversely proportional to the cell radius (or, linearly related to SA/V), because nutrient uptake rate is constrained by surface area, while cellular nutrient quotas are constrained by volume. One assumption that is challenged for larger microbes is constant intracellular nutrient concentration. For example, several microbial species have adaptations like vacuoles that decrease intracellular nutrient concentrations (Menden-Deuer and Lessard 2000).

Combining Equations 1 and 3, we make a prediction for the ribosomal protein mass fraction required to fulfill a change in specific nutrient uptake rate with size (where *max*(*NU*) denotes the nutrient uptake constraint),

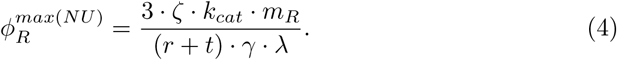

By connecting the simple geometric constraints on specific nutrient uptake with ribosomal investment, we assume that the necessary change in nutrient uptake rate with size will be matched by *sufficient* translating ribosomes. We predict a decline in ribosomal protein mass fraction with increasing cell size under nutrient limitation, 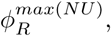 depicted for spherical cells with constant internal nutrient concentrations by the dashed line in Fig. 1.

We are mechanistically modelling two biophysical constraints on maximum growth rate which lead to two associated constraints on ribosomal mass fraction: space limitation (solid line, Fig. 1C) and nutrient limitation (dashed line, Fig. 1C). For a given cell size, the minimum of the two ribosomal protein mass fractions represents the controlling factor. The model requires two free parameters (*k_cat_* for nutrient transporter kinetics in Equation 4, and *ϕ_Q_* the maintenance proteome proportion in Equation 2), which we fitted to empirical ribosomal protein mass fraction data (see below), but these parameter values do not change the core model predictions.

There are three core predictions from this model. The first is that ribosomal protein mass fraction (and maximum potential growth rate) will initially increase from the smallest organisms to some intermediate value. The second prediction is that after this initial increase, ribosomal protein mass fraction (and maximum potential growth rate) will decrease. Third, we predict that the size at which ribosomal protein mass fraction is maximal should correspond to the fastest growing organisms. We also note that at any given cell size, the potential maximum growth rate constrained by space or nutrient uptake will only be achieved by some organisms, while other cell types have evolved for specific environmental conditions where the size-dependent maximum potential growth rate is not achievable or key to fitness.

### 2.2 Ribosomal protein mass fraction as a function of organism volume and maximum growth rate

We aggregated a dataset to test our hypotheses about the controls on maximum growth rate and the associated ribosomal mass fraction in prokaryotes. There are three primary data types we required (Fig. 1B): 1) microbial proteomes (from which we estimated ribosomal protein mass fraction), 2) cell sizes and shapes, and 3) observed growth rates (from which we calculated the maximum observed growth rate). We examined published proteomic data from prokaryotes to test our model (Wang et al. 2012; Kojima et al. 2015; Smith et al. 2016; Müller et al. 2020; Huang et al. 2023; Hou et al. 2023; Zhu et al. 2025), and developed a generalizable statistical approach to adjust for a systematic bias we discovered due to variation in the number of proteins detected (see Methods). Overall, we aggregated more than 200 bias-corrected ribosomal protein mass fractions. For every species, we wanted to identify the proteome where the microbe was likely growing its fastest, so we took the maximum ribosomal protein mass fraction across culture conditions resulting in 97 unique values (one per species). We then determined a range of size estimates for every species to calculate their corresponding volumes using published literature (see Methods).

We identified an increasing pattern in ribosomal protein mass fraction as a function of organism volume, followed by a subsequent decrease (Fig. 1C), mirroring the observed nonlinear pattern in maximum growth rate with cell size (Lynch et al. 2022). Specifically, to test the model we needed to identify all taxa lying along the boundary of the distribution of the observed ribosomal protein mass fraction. We therefore calculated the maximal ribosomal protein mass fractions within logarithmically binned cell volumes to test our model, across varying numbers of bins (Fig. 1C; see Methods), which allowed us to categorize organisms lying on this boundary (coloured in red in Fig. 1C). We examined only the maximal cases because our model makes predictions regarding the limits of fast growth (the *potential* maximum ribosomal protein mass fraction), and there are many reasons why organisms might have slower growth, for example due to suboptimal culturing conditions or evolutionary forces which select for other traits. In the raw data we observed an initial increase, a maximum around 1 *µm*^3^, and a subsequent decrease (Fig. 1C), which is qualitatively consistent with our model. From our fitted model, the size at which ribosomal protein mass fraction was maximized is similar to the fastest growing microbes (*∼*1-10 *µm*^3^; Lynch et al. 2022; Westoby et al. 2021), even though the model was not fitted to any growth rate data. Finally, we also separated the larger prokaryotes and directly correlated maximum observed growth rate with the SA/V, resulting in a Spearman correlation coefficient of 0.63 (we considered this relationship because the nutrient uptake constraint is strictly a model of SA/V, while the results in Fig. 1C are shown as a function of volume; Supplementary Fig. S2).

The model (Equation 1) assumes a relationship between ribosomal protein mass fraction and maximum growth rate which we tested by aggregating previously published growth rate data for all taxa categorized as lying along the boundary of the distribution of ribosomal protein mass fractions (Fig. 1C). We found that ribosomal investment and the maximum observed growth rate are positively correlated (Spearman correlation coefficient = 0.76; Fig. 1D). Our initial translation elongation rate parameter was taken from experiments on *E. coli*, which is used in the equation connecting growth rate with ribosomal protein mass fraction (Equation 1; the solid line in Fig. 1D). For *E. coli*, the predicted maximum growth rate (2.49 *hour^−^*^1^) was similar to the observed maximum growth rate (2.08 *hour^−^*^1^) (Fig. 1D). Similarly, when using recently published estimates of the maximum translation elongation rate (21 amino acids *s^−^*^1^ ribosome*^−^*^1^) for *Vibrio natriegens* (Zhu et al. 2025), we also observe a close correspondence with our predicted (2.53 *hour^−^*^1^) and observed (2.66 *hour^−^*^1^) maximum growth rate (Fig. 1D).

Many organisms have slower maximum growth rates compared with their ribosomal protein investment when using *E. coli* maximal translation elongation rates (Fig. 1D; many points are above the solid line). We speculate that these organisms have slower maximum translation elongation rates or a higher proportion of inactive ribosomes (a rate of 5 amino acids *s^−^*^1^ ribosome*^−^*^1^ is shown on Fig. 1D to illustrate the effect of varying translation elongation rate).

Overall, the two biophysical constraints we describe predict the biggest impact on maximum growth rate and ribosomal allocation for the smallest and largest prokaryotes (Fig. 1C), which we also observe in data (Fig. 1D). This observation led us to examine the consequences of our model for the giant and ultrasmall prokaryotes.

### 2.3 Connecting low ribosomal investment in giant prokaryotes to their metabolism

Our results suggest that as prokaryotic cells greater than *∼* 1*µm*^3^ increase in size, geometric constraints (i.e., a decreasing SA/V ratio) lower their maximum growth rate, in turn leading to a decrease in the necessary ribosomal protein mass fraction (Fig. 1). This decrease potentially frees up proteomic resources for other processes. We therefore asked which cellular processes that require a significant fraction of protein are enriched in low SA/V prokaryotes?

A previous review paper on ‘giant’ bacteria (operationally defined as those with a linear dimension greater than 10 *µm*) noted, qualitatively, that it is “striking that many of the particularly large prokaryotes are either cyanobacteria or sulfide oxidizers” (Schulz and Jørgensen 2001). This led us to hypothesize that the proteins required for these energy metabolisms offset the reduced ribosomal investment in these larger bacteria. To address this hypothesis we aggregated a dataset of known giant prokaryotes and classified the metabolism of individual species (Methods). To compare the metabolisms of giant bacteria with a set of non-giant prokaryotes, we leveraged a published prokaryotic trait database (Madin et al. 2020). Then, we estimated the probability of observing a metabolism in both giant and non-giant prokaryotes by determining a metabolism enrichment value (the ratio of estimated probabilities for giant prokaryotes vs. other prokaryotes). A high enrichment value (i.e., greater than one) indicates that it is more likely that a giant bacterium harbours a given metabolism. We found an estimated enrichment value of 17.4 (8.63-34.0, 95% credible intervals) for “Oxygenic photoautotrophy” and 8.76 (5.57-13.2, 95% credible intervals) for “Oxidation of sulfide, sulfur, thiosulfate” (Fig. 2A). These results complement similar conclusions about phototrophy (Westoby et al. 2021) and suggest that energy metabolisms, like oxygenic photosynthesis and sulfur oxidation, are enriched in giant prokaryotes. But, we were still uncertain if these two metabolisms required a significant protein investment.

**Fig. 2.**
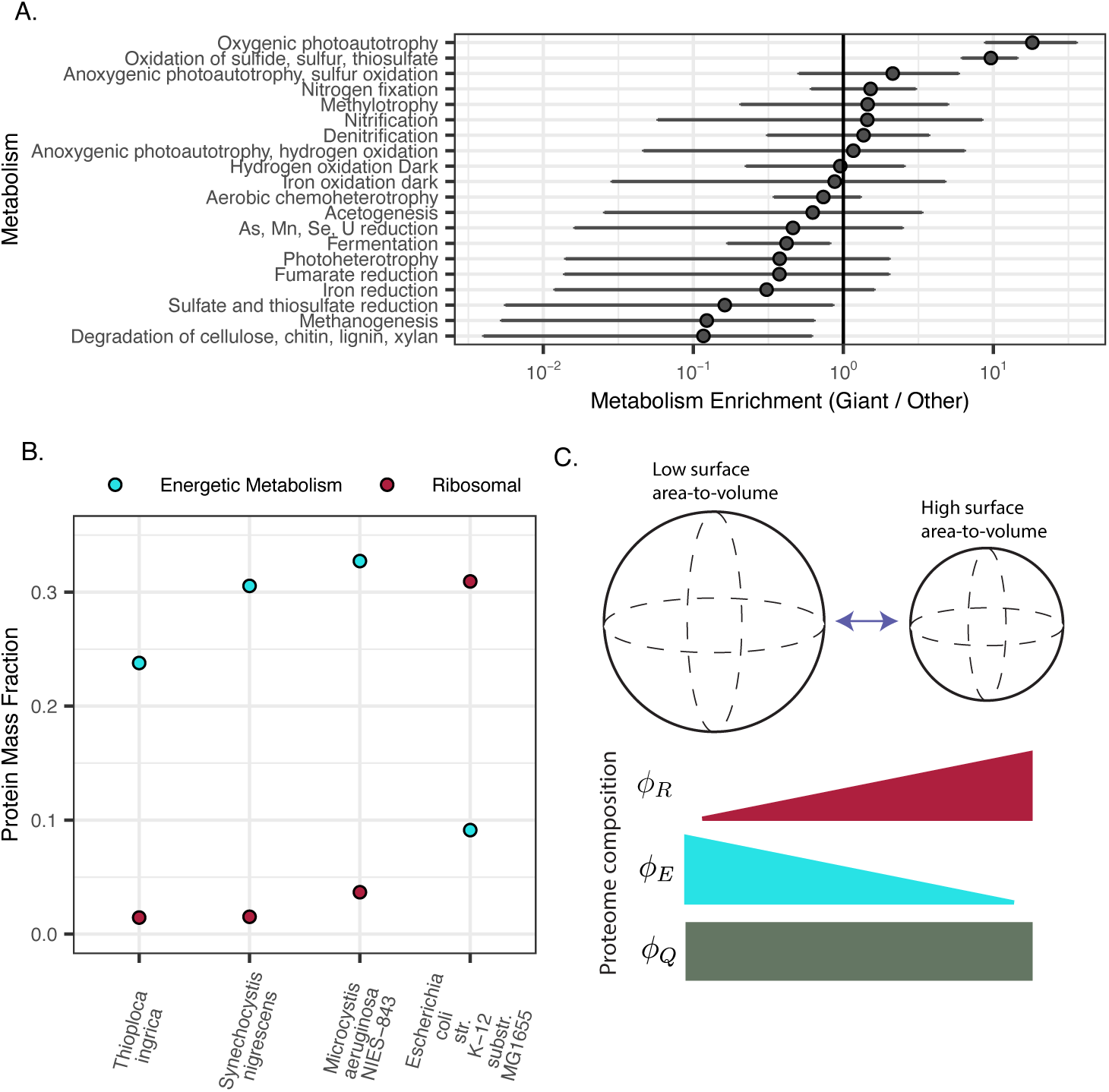
Enrichment in expensive energy metabolisms for low surface area-to-volume prokaryotes. (A) The metabolism enrichment of prokaryotic metabolism, comparing giant prokaryotes vs. a database of prokaryotes. Metabolism enrichment is defined as follows: first, we estimated the probability of a set of prokaryotes as having the ability to conduct a given metabolism. Second, we divided the estimated probabilities from the giant by the other prokaryotes. (B) Protein mass fractions for different sectors for three large prokaryotes. *E. coli* is also shown. (C) A conceptual model predicting that low SA/V prokaryotes (larger sphere) have lower ribosomal protein mass fraction (*ϕ_R_*), which enables a larger investment in energy metabolism proteins (*ϕ_E_*), assuming a fixed, maintenance protein mass fraction (*ϕ_Q_*).

To evaluate whether energy metabolism proteins are a significant proportion of protein mass in large cells (i.e., comparable magnitude to ribosomes in fast growing cells), we quantified investment in proteins involved in energy metabolism for the large prokaryotes in our proteome and cell size dataset, which includes two phototrophs (*Synechocystis nigrescens* and *Microcystis aeruginosa*), and one sulfur oxidizing bacterium (*Thioploca ingrica*). We quantified the proteomic investment in photosynthesis and in sulfur oxidation by categorizing proteins that belong to the Database of Clusters of Orthologous Genes (COGs) category “Energy production and conversion” (using eggNOGG, Herńandez-Plaza et al. 2023). Our results indicate that while the ribosomal mass fraction is less than 5% across giant bacteria, photosynthetic proteins account for *∼*30-33% of total protein in the large phototrophs in our dataset, while sulfur oxidation proteins account for *∼*24% of total protein in *Thioploca ingrica* (Fig. 2B; Kojima et al. (2015)). These results mirror previous work that found large investments in photosynthetic proteins in phototrophs (Jahn et al. 2018; Zavřel et al. 2019; McCain et al. 2022; Burnap 2015). These results are also consistent with the idea that the energy metabolisms of giant bacteria take up a large fraction of their protein biomass, and that this is made possible by a relatively lower investment in ribosomal proteins (Fig. 2C).

A comparison with *E. coli* as a model fast-growing species shows a reversed trend in investment in ribosomal proteins vs. energy metabolism proteins, illustrating this energy-harvesting and growth rate trade-off (Fig. 2B). Further, the SA/V difference between *E. coli* and the three aforementioned large bacteria (Fig. 2B) is relatively large, considering the distribution of SA/V across prokaryotes (see Supplementary Fig. S3 and Methods). Taken together, our findings suggest that at low SA/V, there is an “opportunity” to use these energy sources that require large proteomic investment, providing a mechanistic connection between energy metabolism and cell size.

### 2.4 The volumetric protein burden in ultrasmall prokaryotes

At vanishingly small cell sizes, we found evidence of a strong volumetric constraint on ribosomes per cell and therefore maximum growth rate (Fig. 1A, C). This led us to conduct a sensitivity analysis of the volumetric constraint model (Equations 1 and 2) for ultrasmall prokaryotes, to quantify various cellular components that most impact maximum growth rate (see Supplementary Methods). This analysis revealed that the maintenance proteome, represented by a scalable fraction of total protein (*ϕ_Q_* in Equation 2), has a dramatic impact on maximum growth rate (Supplementary Fig. S4). Increased investment in maintenance proteins takes away from the volume available for ribosomes, and therefore *decreases* maximum growth rate (Equations 1 and 2). The smallest prokaryotes are most impacted because the maintenance proteome parameter is a proportion of *total* (envelope and cytosolic) protein, and at smaller sizes, there is proportionally more envelope protein. Interestingly, we found that the genome volume itself only has a minor impact on growth rate even at very small sizes (Supplementary Fig. S4). In contrast, the volume taken up by proteins could have a large negative impact on maximum growth rate (Supplementary Fig. S4). These analyses are corroborated by measurements in the engineered ‘minimal cell’ (Gilbert et al. 2021; Thornburg et al. 2026), and lead to two hypotheses for specific adaptations in ultrasmall cells that could ameliorate the volumetric constraint (Fig. 3A, B):

1. Ultrasmall prokaryotes have smaller highly expressed, non-ribosomal proteins.
2. Ultrasmall prokaryotes have smaller membrane proteins.

**Fig. 3.**
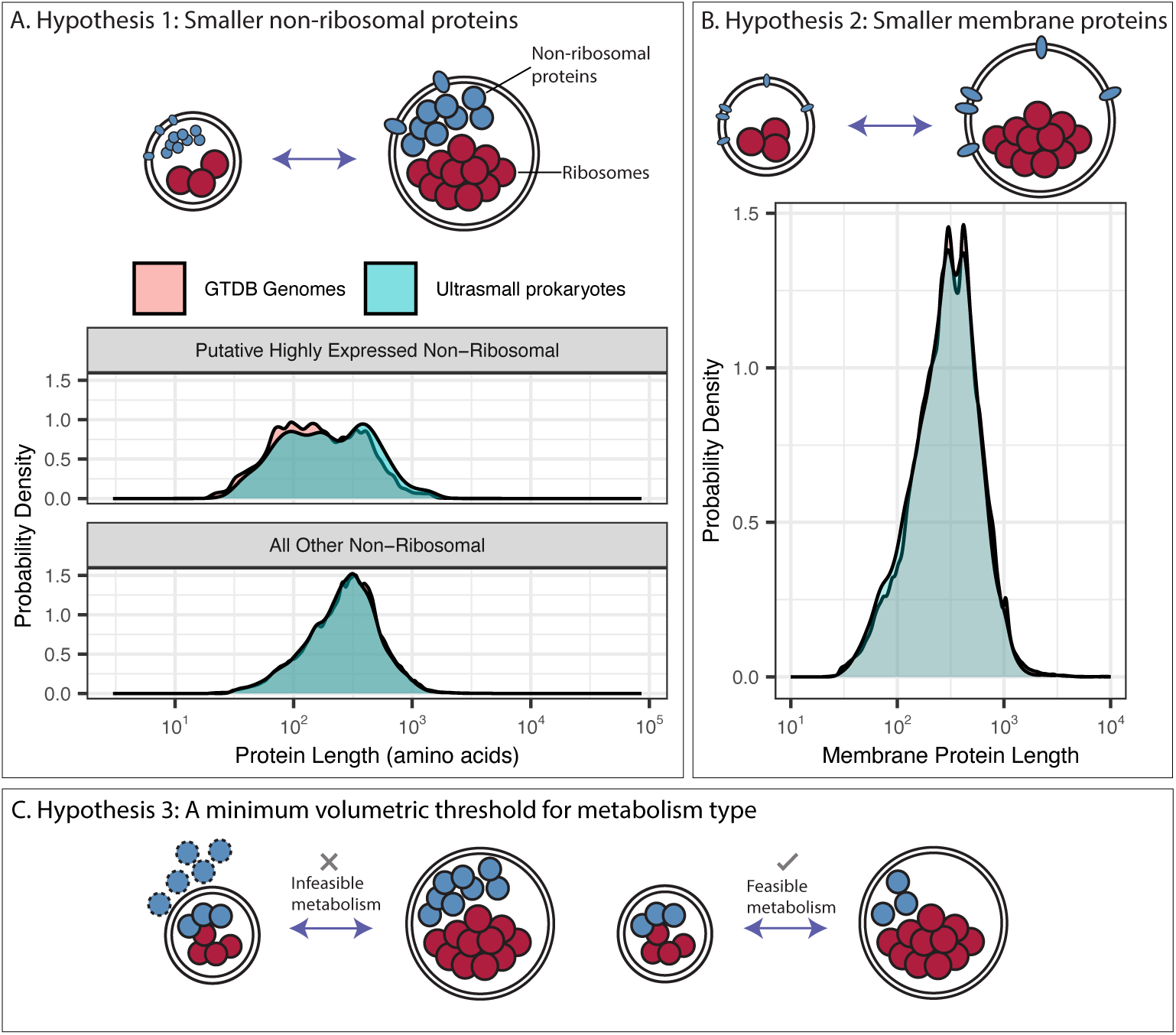
Hypotheses for adaptations in ultrasmall prokaryotes for minimizing the volumetric burden of proteins. Red circles represent ribosome particles and blue circles represent non-ribosomal proteins (not to scale). (A) Hypothesis 1: Ultrasmall prokaryotes evolved smaller non-ribosomal proteins. Density plots comparing the protein length of ultrasmall prokaryotes and a subset (one per prokaryotic family) of the Genome Taxonomy Database (GTDB) genomes. Data are subset by which protein coding genes have putative highly expressed non-ribosomal proteins. (B) Hypothesis 2: Ultrasmall prokaryotes have smaller membrane proteins. Densities for the lengths of membrane proteins for ultrasmall prokaryotes and GTDB genomes. (C) Hypothesis 3: there exists a minimum volumetric threshold for a metabolism type.

To test our hypotheses, we curated a dataset of published ultrasmall prokaryotic genomes (21 individual genomes) and compared them to a reference database across the prokaryotic tree of life (3834 individual genomes, with one genome randomly selected per prokaryotic family from the Genome Taxonomy Database; Parks et al. 2022). The distributions of total protein coding genes were similar across these two groups (Supplementary Fig. S5), further supporting our conclusion that the genome volume is not likely to constrain growth for many cells (Supplementary Fig. S4).

We examined the length distribution across putative highly expressed, non-ribosomal proteins (expression level predictions were made using codon bias, see Methods; Supek and Vlahovčcek 2005), and across all other non-ribosomal proteins. We found no evidence for a difference in non-ribosomal protein lengths comparing ultrasmall prokaryotes and other prokaryotes, regardless of the predicted expression level (Fig. 3A). Next we compared the length of membrane proteins in ultrasmall prokaryotes versus the GTDB subset, and again found no evidence that membrane proteins are systematically smaller in ultrasmall prokaryotes. A key unknown is the expression level of each protein, which could be systematically different across size classes. While we observe no differences in the length of highly expressed or membrane proteins in ultrasmall cells, we noticed that all appear to be heterotrophs (although the designation of metabolism in these poorly characterized species is challenging). This observation led us to our third hypothesis (Fig. 3C):

1. Ultrasmall prokaryotes use metabolisms with an intrinsically low minimal protein volume threshold.

There may be a *minimal* volumetric protein burden associated with any given metabolism. Autotrophy could be an infeasible metabolism because it has a higher intrinsic minimum protein investment compared to heterotrophy (Fig. 3C). Each of these metabolisms requires a set of proteins, which we posit varies in the minimal required contribution both in terms of number of unique proteins and their corresponding copy number (the required copy number is the largest source of uncertainty in our hypothesis; see Supplementary Fig. S6).

## 3 Conclusions

We put forward a mechanistic model for two phenomena: 1) the initial increase and subsequent decrease in maximum growth rate with organism size, and 2) a corresponding proteomic signature, the ribosomal protein mass fraction. By aggregating a large dataset of proteomic data from a diverse set of prokaryotes, we find empirical support for two mechanisms constraining maximum growth rate (Fig. 1A). We also find that changes in ribosomal protein mass fraction are a central factor in shaping the metabolisms of giant and ultrasmall prokaryotes.

Our framework may be useful for identifying microbes that have unique translation properties like elongation rate or other cellular properties. One intriguing bacterium in this dataset is *Caldisericum exile* (Mori et al. 2009), an anaerobic thermophilic bacterium discovered in 2009, which our analyses revealed to have many ribosomal-like proteins that have low sequence homology to other prokaryotes. We speculate that its ribosomal structure is significantly diverged from other microbial taxa, making it an exciting candidate to explore atypical translation mechanisms.

The two constraints on maximum growth rate we examined had the strongest predictions for the very large and very small prokaryotes, and we connected these constraints to metabolism of these size classes. But, there is a large swath of prokaryotic sizes in the middle. What explains the diversity of metabolisms within this middle group? We speculate that ecological trade-offs in resource availability may govern the elevated metabolic diversity in this size class. For example, a 1 *µm*^3^ phototroph will likely grow slower than its heterotrophic counterpart due to higher protein cost of phototrophic metabolism (Burnap 2015), but it would have the ecological advantage of using light for growth (i.e., maximum growth rate is only one of many important ecological dimensions; Weissman et al. 2021; Zakem et al. 2024; Hoffert et al. 2026).

There are several extensions of our work. First, the empirical relationship between maximum growth rate and cell volume is used extensively in ecosystem-scale models (Follows et al. 2007; Dutkiewicz et al. 2009). Mechanisms that underpin this relationship are particularly important for parameterizing how environmental factors, like temperature, will change maximum growth rate. Temperature could be explicitly parameterized within our model, where kinetic processes might have different temperature dependencies (e.g., translation vs. nutrient uptake), and therefore produce different outcomes for the scaling of maximum growth rate and organism size.

A major extension will be applying our model to eukaryotes, which could be parameterized given the differences in ribosome-particle mass and maximum translation elongation rates. Recent work in mammalian cell lines has provided evidence consistent with the connection between surface area, cell volume, and growth rate (Mu et al. 2020; Wu et al. 2025). But, this work also highlights that applying our framework to eukaryotes may be challenged by accurately quantifying surface area (Wu et al. 2025). Broadly, it seems likely that the mechanistic constraints on maximum growth rate change across the eukaryotic size spectrum (Bozdag et al. 2023; Banavar et al. 2002; DeLong et al. 2010; Hoehler et al. 2023).

### 3.1 Limitations of the study

One persistent challenge in our analyses is uneven data density across organism sizes. We searched for additional proteomic datasets at the size extremes to fill in those gaps. For example, we used proteomic data from the giant bacterium *Thioploca ingrica* (Kojima et al. 2015). More data are required to further test our model, particularly for extremely small or large bacteria. Another limitation of our analysis is the lack of simultaneously collected growth rate, proteomic, and size data. This was a necessary trade-off to maximize the diversity of microbes we included in our analysis, but, ideally, these three components would be collected in tandem. Finally, identifying an intrinsic maximum growth rate for an organism is challenging, and typically requires growing it under numerous conditions to identify an optimum.

We used several approaches for comparing microbial genomes or metabolism across size classes (e.g., to examine the potential enrichment of certain metabolisms in giant bacteria, or the protein length distribution in ultrasmall prokaryotes). A challenge with quantifying enrichment is the requirement to define a baseline, which inevitably has several biases. For example, the high prevalence of fermentation in the phenotypic trait database of bacteria and archaea (Madin et al. 2020) is likely biased towards: 1) microbes with metabolisms that can easily be grown. 2) Metabolisms that are simpler to measure, and 3) highly-studied microbial taxa like pathogens. It is unclear how to adjust for these types of biases.

Trickovic and Lynch (2025) proposed that there is an increased requirement for lipid biosynthetic machinery for very small organisms, as lipids are an increasingly large portion of cell mass as cell size decreases. We have not ruled out their model, but the model proposed above is an alternative (and arguably simpler) explanation. In both of our analyses, we identified an increase in the maximum ribosomal protein mass fraction with volume for small prokaryotes (albeit with overlapping data sources). In other problems, distinguishing between space constraints and protein costs has been challenging because these two constraints are sometimes unidentifiable (Basan et al. 2015). We suspect that experimentation with new model organisms, for example, the very small bacterium *Mesoplasma florum* (Matteau et al. 2020), will help to distinguish between these mechanistic models.

Finally, we note that it is plausible that there was natural selection for a specific metabolism, like sulfur oxidation, and subsequently these organisms evolved lower SA/V, or vice versa. Our model is not able to distinguish the stronger selected effect. Alternative hypotheses regarding the additional structures associated with these metabolisms, e.g., thylakoid membrane space, also remain to be quantitatively examined.

## 4 Methods

### 4.1 Quantifying ribosomal protein mass fraction across diverse prokaryotes

We examined published proteomic data from prokaryotes to test the model (Wang et al. 2012; Kojima et al. 2015; Smith et al. 2016; Müller et al. 2020; Huang et al. 2023; Hou et al. 2023; Zhu et al. 2025). The first step is to uniformly annotate proteins as ribosomal or not, and the second step is to accurately quantify ribosomal protein mass fraction.

We identified ribosomal proteins using trained profile Hidden Markov Models (HMMs) for Pfams that correspond to ribosomal proteins (see list of Pfam IDs in Supplementary Table S2). To identify the set of Pfams to include, we first searched the Pfam database with the string “ribosom”. We then manually curated the list of resulting Pfams to identify structural components of the ribosome and gathered their associated IDs (Supplementary Table S2). We then classified all protein sequences as ribosomal or not from the protein sequence file used for database searching for the discovery proteomics of these organisms. We used an E-value cutoff of 10*^−^*^2^, and our ribosomal protein mass fraction estimates are largely insensitive to this choice (Supplementary Fig. S7).

To calculate ribosomal protein mass fraction for data from PaxDb (Wang et al. 2012; Müller et al. 2020; Huang et al. 2023), we needed to convert the protein abundances (in ppm) into quantities proportional to mass fraction. We therefore multiplied protein abundances by their corresponding protein length (also derived from the fasta file used for database searching). We summed these transformed (weighted by protein length) protein abundances for each organism, which we used as an organism-specific normalization factor. Each transformed protein abundance, for each organism, was then divided by this normalization factor. Finally, we summed all the normalized ribosomal protein abundances to calculate a species-specific ribosomal protein mass fraction.

We used several additional datasets that required different methods for calculating ribosomal protein mass fraction. To calculate ribosomal protein mass fraction for data from *Thioploca ingrica* (Kojima et al. 2015), we followed Ishihama et al. (2005) for calculating the protein mass fraction (Equation 4 in Ishihama et al. (2005)). To calculate ribosomal protein mass fraction for data from *SAR11* proteomes (Smith et al. 2016), we first normalized all peptide abundances by the sum of identified peptide abundances. We then summed up all normalized, ribosomal peptide abundances to estimate the ribosomal protein mass fraction. For the *Synechocystis nigrescens* proteomic data (Müller et al. 2020), we first normalized all ribosomal protein intensities by the sum of all identified protein intensities. We then summed up all ribosomal proteins, which approximates the mass fraction (Cox et al. 2014). In each of the above cases, we annotated all protein coding genes as ribosomal or not (the same fasta files that were used for database searching) using profile HMMs as above.

Previous work using mass spectrometry simulations has shown that estimating proteomic composition can be biased when the number of proteins (or peptides) detected changes (McCain and Bertrand 2019; McCain et al. 2022). We empirically examined this phenomenon by leveraging a well studied bias in mass spectrometry, where abundant proteins are more likely to be identified. An abundant-protein bias is explicitly encoded in many mass spectrometry experiments, where the most abundant peptides are preferentially selected (“TopN” ion selection in data-dependent acquisition). Specifically, we subsetted our dataset of 97 proteomes to only those with more than 1000 proteins identified. Then, we subsampled the top *n* most abundant proteins (where *n* = 400, 500, 700, or 1000). After subsampling, we calculated the ribosomal protein mass fraction, and compared the subsampled ribosomal protein mass fraction with the ribosomal protein mass fraction with all observed proteins. These simulations showed that as the total number of proteins identified decreases, the ribosomal protein mass fraction tends to be overestimated. It also showed that higher ribosomal protein mass fractions are more sensitive to overestimation than lower ribosomal protein mass fractions (Supplementary Fig. S8).

We used these simulations to parameterize a simple correction method, where the inputs are the total number of proteins identified (*P*) and the observed ribosomal protein mass fraction (*ϕ^Obs^*). Specifically, we optimized *α* to predict the true ribosomal protein mass fraction (*ϕ_R_*) as a function of the above inputs and one fitted parameter,

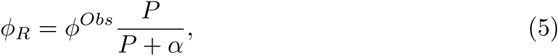

with the fitted value of *α* = 62 identified proteins (Supplementary Fig. S9 shows the observed and bias-corrected ribosomal protein mass fractions per species). We suspect this bias correction method would be useful in metaproteomics, where the abundance of organisms (and therefore the number of proteins detected) can vary widely.

### 4.2 Microbial volume calculations

We searched for documented dimensions, shapes, and/or volumes, for prokaryotic microbes that also have a published proteome. We unfortunately did not have cell volumes or dimensions for the corresponding proteomes. We therefore collected ranges of dimensions for different taxa where possible. In most cases, we used reported ranges of all given dimensions, and calculated the minimum and maximum volumes, assuming the dimensions are independent. For each organism, we classified shapes as spherical, cylindrical, or box-like. In some cases, we found published values on volumes and we then used the volumes directly. In cases where numerical values were not reported in the literature, we used microscopy images and ImageJ to quantify the maximum and minimum dimensions for individuals within the field of view. In these cases where we had access to raw images, we did not assume dimensions were independent. All recorded measurements and sources are detailed in Supplementary Dataset S1.

### 4.3 Aggregating microbial growth rates

We collected previously published growth rate data from the subset of microbes that lie along the border of maximum ribosomal protein mass fraction across sizes (Fig. 1C). We restricted our analysis to this subset for two reasons: 1) we are trying to identify the experimental conditions where ribosomal protein mass fraction was maximized within an organism, and it is possible that the organism was only cultured in a single (slow growing) condition. 2) There are many reasons why an organism within a size class has a lower maximum growth rate. For example, the maximum phototrophic growth rate is much slower than the maximum heterotrophic growth rate (Burnap 2015). Further, maximum growth rate is only one ecological dimension, and we suspect that some variation in the maximum growth rate *within a size class* is a function of metabolism. Observed growth rates were aggregated from diverse published sources (Madin et al. 2020; Scott et al. 2010; Mateus 2025; Buyanbadrakh et al. 2026; Evsyutina et al. 2022; Noecker et al. 2023; Ng and Hamilton 1971; Matteau et al. 2020; Schaus et al. 2024; Jones and D’Orazio 2013; Budinich et al. 2011; Williams et al. 2024; Dai and Koehler 1997; Yang and Miller 2008; Passalacqua et al. 2007; Feng et al. 2025; Aranda-Díaz et al. 2020; Wilson et al. 2006; Lankiewicz et al. 2016; Rappé et al. 2002; Weinstock et al. 2016; Adler 2015; Van Alphen et al. 2018; Abboud et al. 2005; Van Der Kooij et al. 2016; Ze et al. 2012; Perez-Sepulveda and Hinton 2025; Alessi et al. 2020), and we subsequently calculated the maximum observed growth rate for each species. For many of these growth rate observations, we identified growth rates that are observed in controlled cultures for the exact species and strains that correspond to the proteomic data. There were two notable exceptions. *Thioploca ingrica*, a proteome we included, was from cells grown in natural environments. To our knowledge these giant bacteria have not been successfully isolated in culture. Our growth rate estimates are crude for this species, coming from early work on hay-enrichment cultures of *Thioploca ingrica* (Maier 1980). A second exception was for the cyanobacterium *Synechocystis nigrescens*. We were unable to find growth rate data for this large cyanobacterium. We used growth rate data from its congener, *Synechocystis* sp. PCC 6803, which is likely an overestimate because it is a smaller cyanobacterium.

### 4.4 Parameterizing and fitting the biophysical model

A full table of parameters identified for our biophysical model is available in Supplementary Table S1 (including translation elongation rates, cytosolic and membrane protein densities, protein mass per ribosome particle, etc.). Note that many of these parameters are derived from model organisms, in particular *E. coli*.

We fitted two parameters to proteomic data: the maximum turnover rate per transporter (*k_cat_*), and the proportion of total protein that is required for cell maintenance (*ϕ_Q_*). To fit the model, we first logarithmically binned the bias-corrected ribosomal protein mass fractions across all our data. We then identified the maximum of each bin, and used *L-BFGS-B* optimization to find parameters that minimized the sum of the absolute value of the differences between the observed ribosomal protein mass fractions and model predictions. Both parameters were constrained with a lower bound of 0, and *ϕ_Q_* was constrained with an upper bound of 1. Initial values were 40 and 0.2 for *k_cat_* and *ϕ_Q_*, respectively. We repeated this optimization procedure using variable numbers of organism volume bins (from 15 to 20), where the minimum and maximum bin values are 0.01 and 100, respectively.

### 4.5 Annotating energetic metabolism proteins

We annotated proteins that contribute to energetic metabolism using eggNOG mapper (Abueg et al. 2024; Herńandez-Plaza et al. 2023). We used the annotations from the Database of Orthologous Clusters (COG) to designate if a protein is part of energetic metabolism, specifically category “C - Energy production and conversion”. One central challenge of using this approach, particularly for comparing across metabolism types, is that the automated annotation may have differential success in labelling proteins *required* for a given metabolism. For example, pigment proteins are required to make pigments used for photosynthesis, but they are not considered as part of the COG Category in Energy production and conversion.

### 4.6 Estimating metabolic enrichment in giant prokaryotes

We aggregated a list of giant bacteria from several previous review papers on these organisms (Schulz and Jørgensen 2001; Ionescu et al. 2023). We then systematically examined previous literature on these organisms and classified their metabolism using the same categories as a prokaryotic trait database (Madin et al. 2020). There is uncertainty about the classification of metabolism because of the enigmatic nature of these organisms, so we used a combination of evidence to classify them (environmental context, genomic composition, etc.).

We quantified the enrichment in metabolism by estimating the probability of observing a metabolism within each size class, and then comparing these estimated probabilities (giant vs. all others in the trait database). If there is evidence of a given metabolism we considered that a binary value, and we model the sum of these values within the group (*y*) following a binomial distribution

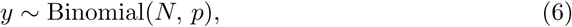

where *N* is the total number of species (either in the set of giant bacteria or the trait database), and *p* is the estimated probability. If all taxa within a size class can ferment, for example, then we would estimate *p* to be close to 1. We then compared the estimated probabilities to obtain a metabolism enrichment value (*E_k_*),

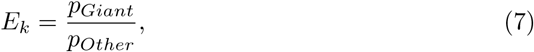

for a given metabolism *k* (shown in Fig. 2A). This statistical model was fitted using Stan and rstan.

### 4.7 Comparing genomes from ultrasmall prokaryotes to GTDB genomes

We compared a curated set of genomes from ultrasmall prokaryotes to a large genomic database of prokaryotes (the Genome Taxonomy Database, GTDB; Parks et al. 2022). We sought to make a balanced sampling of the phylogenetic diversity from GTDB, so we randomly selected one genome from each prokaryotic family, resulting in 3834 genomes in total. For curating the set of ultrasmall prokaryotes, we made a list of named species in several published review papers on these organisms (Duda et al. 2012; Nakai 2020), and manually searched for published genomes.

Prodigal was used to annotate protein coding genes in each genome (both the ultrasmall and GTDB genomes; Hyatt et al. (2010)), and we then annotated ribosomal proteins using profile HMMs as described above, using an E-value cutoff of 10*^−^*^2^. Then, we predicted which proteins are highly expressed using a codon adaptation index, Measure Independent of Length and Composition (MILC; Supek and Vlahovčcek (2005)), with the reference group being the set of annotated ribosomes. There are major assumptions about predicting expression level using a codon adaptation index; however, we do not have access to proteomic or transcriptomic data associated with this large dataset. We used a top 5% cutoff of the MILC expression index to conservatively predict whether or not a gene is highly expressed (empirically informed by previous work; Klumpp et al. (2012)). For identifying membrane proteins, we used TMHMM 2.0 (Krogh et al. 2001), where a protein is considered a membrane-protein if there are at least two predicted membrane helices.

## Data Availability

All data and code are available for download via Figshare (https://figshare.com/account/articles/34112706).

## Acknowledgments

This work benefited from discussions with Christopher Follett and others within the MIT Darwin group. Thank you to Petra Louis for supplying raw growth rate data on *Coprococcus comes*. Thank you to Noor Youssef and Debora Marks for suggestions on the analysis of ultrasmall prokaryotic genomes. Thank you to Qingyao Huang for correspondence about PaxDb. J.S.P.M. is a Damon Runyon Fellow supported by the Damon Runyon Cancer Research Foundation (DRG-2470-22). M.J.F. is very grateful for support from the Simons Foundation (CBIOMES Award 549931).

## Notes

### Competing Interest Statement

The authors have declared no competing interest.

