## Supplementary Methods and Figures for "Proteomes reflect size-based growth constraints across diverse microbes"

### 1 Supplementary Methods

#### 1.1 Trait database for surface area-to-volume ratios

We used a bacterial and archaeal trait database to determine a distribution of SA/V across various prokaryotes (Madin et al. 2020). There are 9 distinct cell shapes recorded in this database, but several of them are challenging to confidently calculate SA/V from (e.g., "pleomorphic", "vibrio"). We therefore considered three categories, "bacillus", "spiral", and "coccus", which comprised around 92% of the entries that contain shape information. There are two dimensions noted in this database, we considered the longer one to refer to the length, and the other to the width.

After categorizing by shape, we estimated volume ($V$) and surface area ($SA$). For all calculations, $r$ represents the cell radius and $L$ represents the total cell length. We treated the cell shapes as follows: Cocci were modeled as a perfect sphere. Bacillus and spiral prokaryotes were modeled as cylinders with hemispherical ends (capsules).

The volume for spherical cells (cocci) follows the standard sphere formula. For bacilli and spiral forms, the volume is calculated as a cylinder of length $(L - 2r)$ with two hemispherical caps of radius $r$, which simplifies to:

$$V = \begin{cases} \frac{4}{3}\pi r^3 & \text{if coccus} \\ \pi r^2 \left(L - \frac{2}{3}r\right) & \text{if bacillus or spiral} \end{cases}$$

The surface area for cocci is defined by the surface of a sphere. For the elongated forms (bacillus and spiral), the surface area includes the side walls of the cylinder plus the area of the two hemispherical caps:

$$SA = \begin{cases} 4\pi r^2 & \text{if coccus} \\ 2\pi r L & \text{if bacillus or spiral} \end{cases}$$

#### 1.2 Volumetric constraint sensitivity analysis

We sought to quantify how various cellular components and processes impact growth rate by conducting a parameter sensitivity analysis on the volumetric constraint. This analysis is done by substituting the volumetrically constrained, ribosomal protein mass fraction ($\phi_R^{max(v)}$) into the expression for growth rate. We then increased every model parameter by 10% and examined the resulting impact on maximum growth rate, across a range of cell volumes.

Several parameters were cell volume-independent while others were volume-dependent. Among volume-independent effects we observed that increasing translation elongation rate increases maximum growth rate, and increasing ribosomal protein mass decreases maximum growth rate, both of which directly follow Equation (1) (where the volume dependence enters separately via $\phi_R^{max(v)}$).

Among volume-dependent effects, we observe that increases in the envelope layer thickness or the envelope protein density lead to similar decreases in maximum growth rate, which results from the need for ribosomes to synthesize relatively more non-ribosomal protein. In contrast, an increase in the density of cytoplasmic protein increases maximum growth rate, especially for the smallest cells, because it relaxes the volumetric constraint on the ribosome mass fraction. These perturbations become stronger at smaller sizes because of the volumetric effect of packing ribosomes is strongest in this size range. The genome volume has a small negative impact on maximum growth rate, and we predict that only at the very smallest cells it can slightly impact maximum growth rate.

#### 2 Supplementary Figures

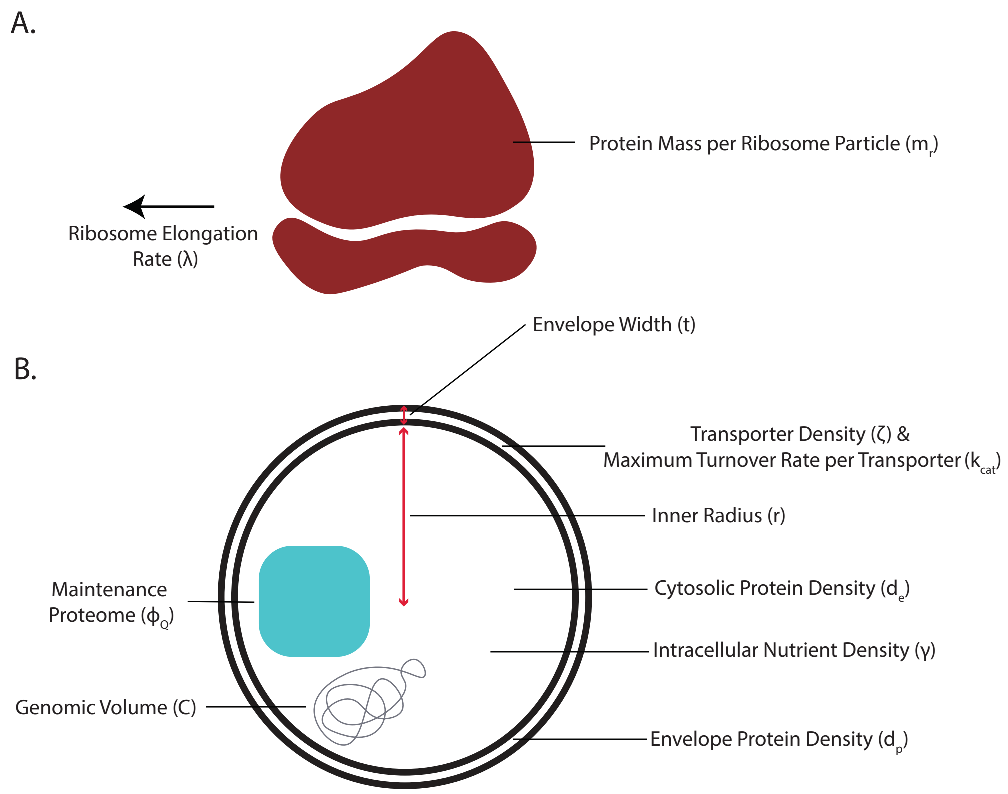


**Fig. S1** Illustration of different parameters for the cellular constraints on maximum growth rate. (A). Parameters related to ribosomes and their corresponding symbols. (B). Parameters related to cell size and nutrient uptake.

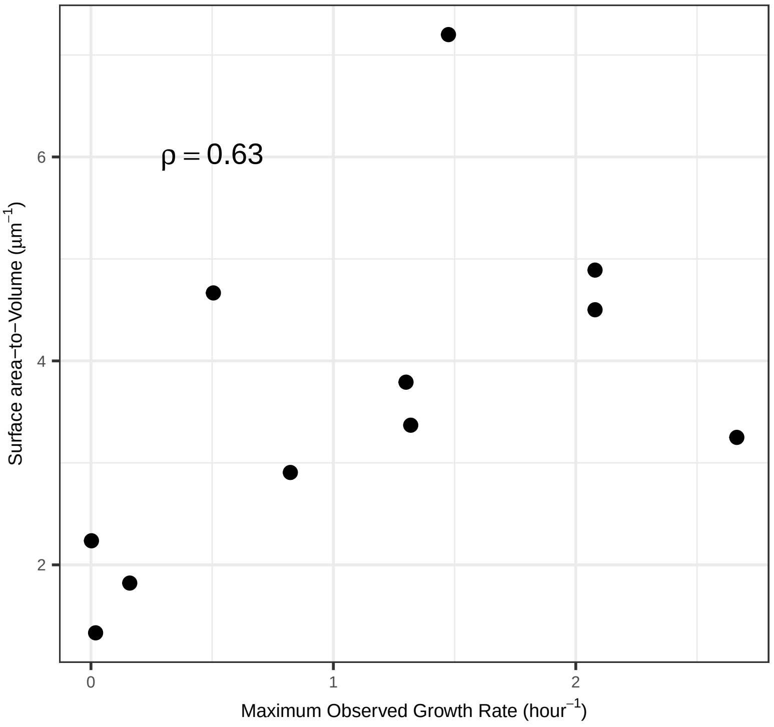


**Fig. S2** The relationship between maximum observed growth rate and the surface area-to-volume ratio for prokaryotes greater than 2 $\mu m^3$.

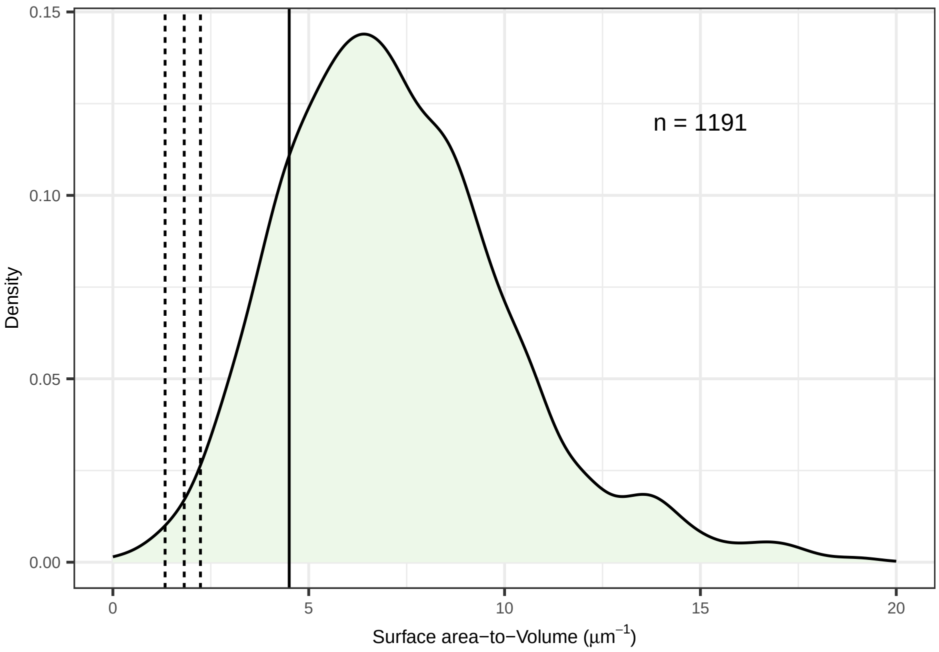


**Fig. S3** The distribution of surface area-to-volume ratios (SA/V) in a published prokaryotic trait database (Madin et al. 2020) shown as a density distribution. The four vertical lines represent the SA/V of different organisms shown in the main text, specifically, the three dashed lines correspond to *Synechocystis nigrescens*, *Microcystis aeruginosa*, and *Thioploca ingrica*. The solid vertical line is the SA/V for *Escherichia coli*.

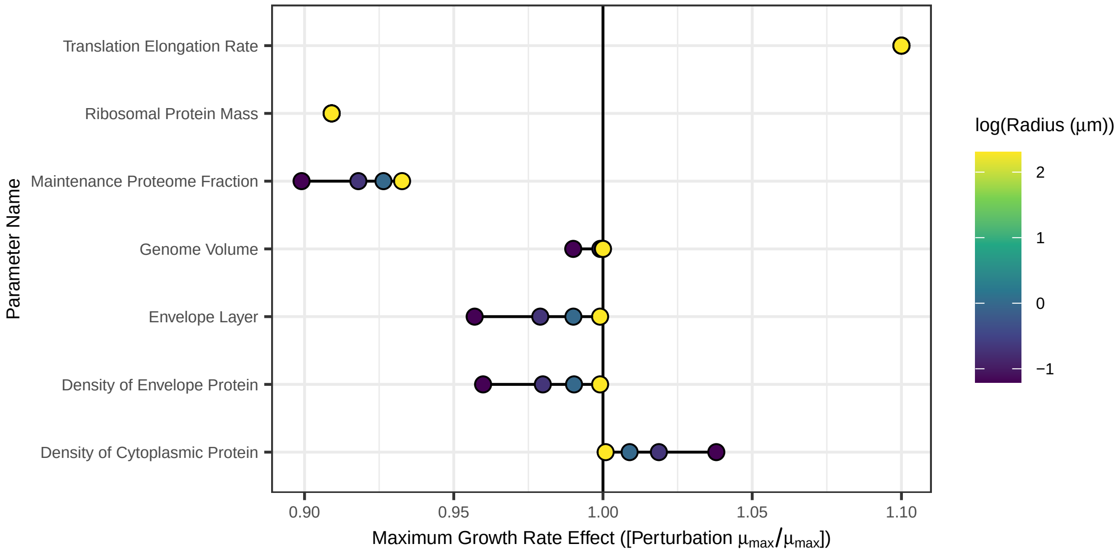


**Fig. S4** Examining processes constraining maximum growth rate at small sizes. (A) Impact on maximum growth rate ($\mu_{max}$) of 10% increases in various processes and cellular properties. The growth rate impact is calculated by dividing the maximum growth rate after perturbation by its the baseline value.

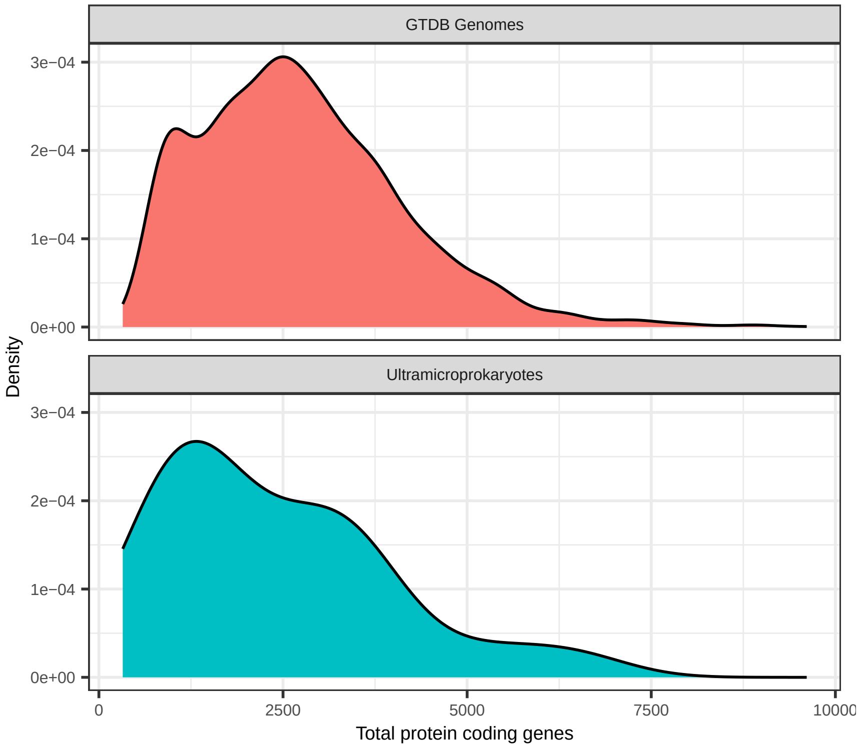


**Fig. S5** Probability densities of total protein coding genes comparing GTDB genomes (3834 genomes) and the set of ultrasmall prokaryotic genomes (21 genomes). The mean (standard deviation) values were 2738 (1379) and 2391 (1537) for GTDB genomes and ultramicroprokaryotic genomes, respectively.

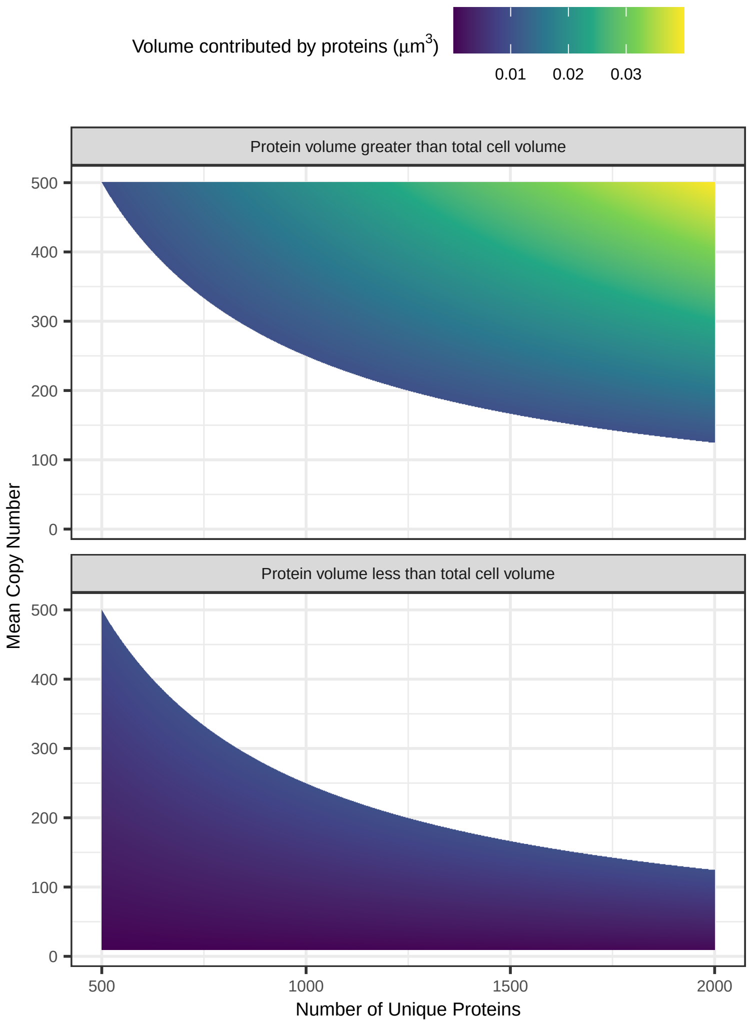


**Fig. S6** The volumetric contribution of proteins across varying degrees of average copy number (every protein is given the same copy number) and number of unique proteins. Volume is calculated assuming an average protein length of 300 amino acids, and 110 Daltons per amino acid, and a conversion factor of 1.212e-3 for moving from Daltons to $nm^3$ (Erickson 2009). The assumed cell volume is 0.01 $\mu m^3$, which distinguishes the top and bottom panel.

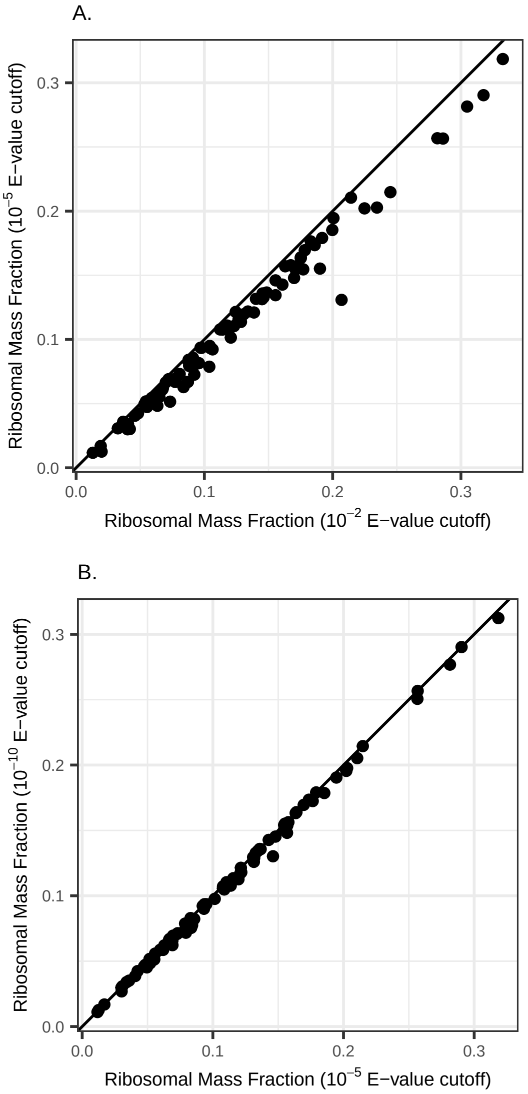


**Fig. S7** Estimated ribosomal mass fractions as a function of different E-value cutoffs. Each point corresponds to the maximum ribosomal mass fraction per species. (A). The horizontal axis shows the ribosomal mass fraction estimated when a $10^{-2}$ E-value cutoff was used, and the vertical axis shows the ribosomal mass fraction estimated when a $10^{-5}$ E-value cutoff was used. (B) The horizontal axis shows the ribosomal mass fraction estimated when a $10^{-5}$ E-value cutoff was used, and the vertical axis shows the ribosomal mass fraction estimated when a $10^{-10}$ E-value cutoff was used. The line in each panel is a 1:1 relationship.

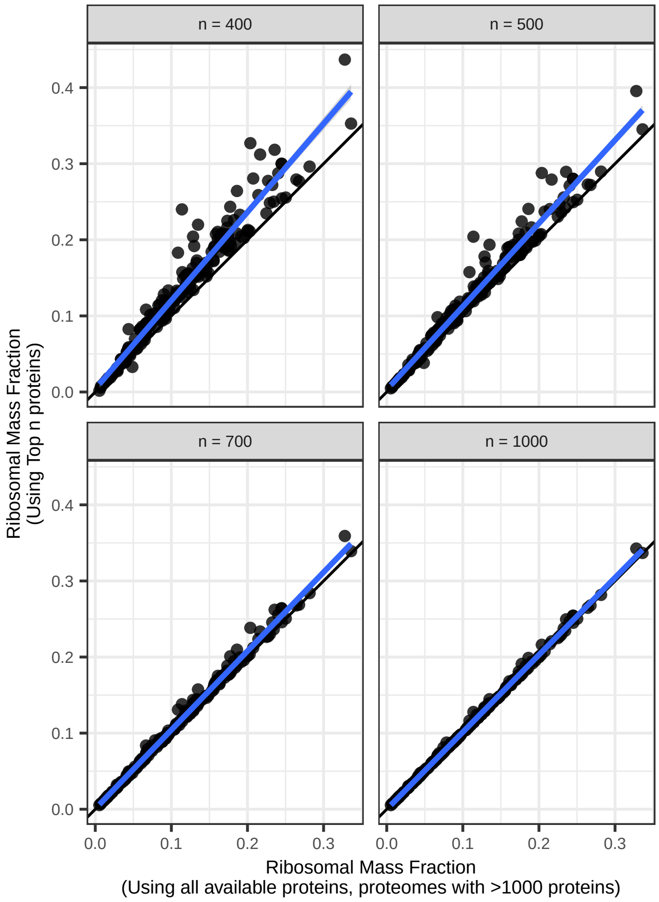


**Fig. S8** Ribosomal mass fraction estimated using the top $n$ most abundant proteins (vertical axes), where different panels correspond to different values of $n$. The horizontal axis is the estimated ribosomal mass fraction using all available proteins for organisms with more than 1000 detected proteins. The lines are 1:1 relationships. Blue lines are linear regressions between top $n$-derived ribosomal protein mass fraction and the all available protein-derived ribosomal protein mass fraction.

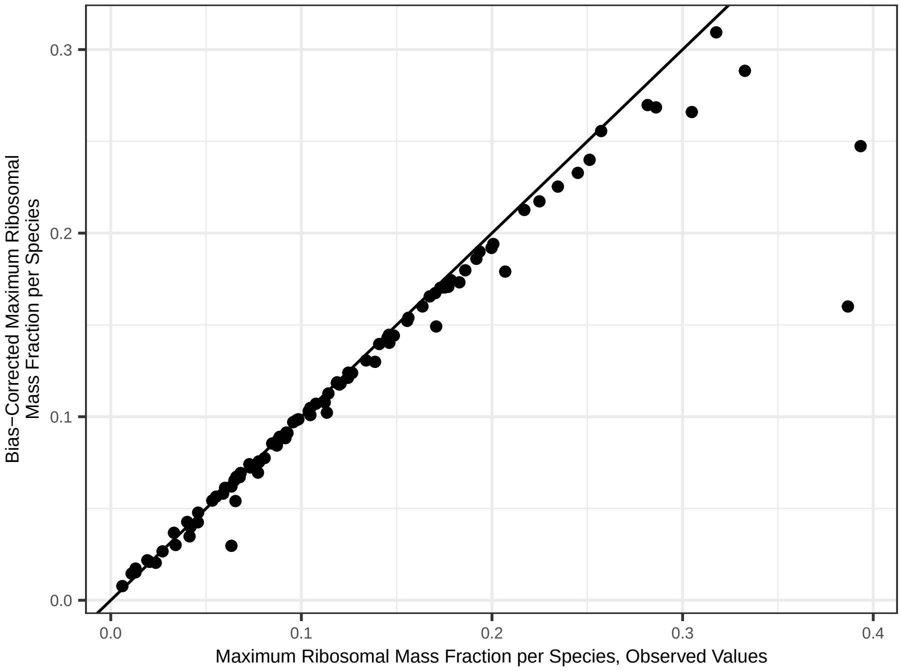


**Fig. S9** The relationship between the observed ribosomal mass fraction (horizontal axis) and the bias-corrected ribosomal mass fraction. Each point corresponds to a maximum ribosomal mass fraction per species. The line is a 1:1 relationship.
